# CRISPR-Cas9 editing efficiency in fission yeast is not limited by homology search and is improved by combining gap-repair with fluoride selection

**DOI:** 10.1101/2024.03.01.582946

**Authors:** Ronan Fernandez, Julien Berro

## Abstract

Protocols for CRISPR-Cas9 editing have been implemented in most model organisms, including fission yeast, for which some improvements have also been later described. Here, we report an improvement to the CRISPR-Cas9 protocol in fission yeast, as we combine a cloning free gap-repair method with our previously described fluoride selection marker, which speeds up genome editing. We also report a wide variability of editing efficiencies at different loci along the genome, and we demonstrate that this variability cannot be explained by the location of the edited sequences in the genome. Lastly, our attempt at improving editing efficiency by targeting the donor DNA to the cut site using a HaloTag strategy to link the donor DNA to two proteins of the homologous recombination repair machinery (Rad51 or Rad52) fell short, which shows that editing efficiency in fission yeast is likely not limited by homology search.

## Introduction

In the past 10 years, Clustered Regularly Interspaced Short Palindromic Repeat (CRISPR) have been repurposed and used in combination with CRISPR-associated (Cas) nucleases as a biological tool for genome editing. CRISPR-Cas9 has since revolutionized the field as a highly competitive alternative to the previously used TALEN (Transcription Activator-Like Effector Nucleases), ZFN (Zinc Finger Nucleases) [1-3] and MN (MegaNucleases) [4, 5] programmable enzymes. Many Cas nucleases with variable properties continue to be identified [6, 7], allowing researchers flexibility according to their organisms of interest, type of genome editing needed and resources available.

However, most Cas9 endonucleases present a high rate of off-target edits [8-10], as well as wide variability in efficiency with respect to the chosen editing site, even sometimes failing at editing some refractory sites [11]. Strategies have been developed to improve Cas9 editing specificity and efficiency including single-guide RNA (sgRNA) modifications [12, 13], using Cas9 nickases [14] and identification of other endonucleases [7]. Recently, engineering strategies such as fusing Cas9 to chromatin-modulating peptides has led to editing efficiency improvement [15], as well as binding the homologous recombination (HR) donor DNA to Cas9 via a monoavidin-biotin interaction [16]. Unfortunately, no universal feature defining a highly efficient Cas9 site or a good repair outcome have been identified so far. Thus, understanding in detail all the steps of CRISPR/Cas9 editing becomes necessary and the first cornerstone to improve efficiency.

CRISPR-Cas9 has been implemented in *Schizosaccharomyces pombe* in 2014 [17] and has tremendously helped the fission yeast community, especially for the fast and markerless construction of strains containing points mutations or N-terminal tagging of genes. We reported in 2016 that fluoride exporter channels (*fex1* and *fex2*) can be repurposed as selection markers in *S. pombe* [18] to significantly improve the speed and efficiency of the CRISPR/Cas9 editing protocol. The description of a ligation free cloning technic based on a PCR-system to create the sgRNA vector [19] shortened the protocol and in 2018, a gap repair method [20] described how to bypass the creation of the gRNA vector, which required DNA cloning, purification and sequencing, the most time-consuming steps of the CRISPR-Cas9 protocols in yeast. Here, we combine the cloning-free gap-repair strategy with our fluoride selection marker to shorten the protocol considerably. We also report a wide variability of CRISPR/Cas9 editing efficiencies along the fission yeast genome and try to tackle the causes of this variability, showing that the genomic location of the attempted editing doesn’t seem to play a major role. We also engineered fission yeast cells that are able to target the donor DNA to the Cas9 cut site but we did not observe any improvement in editing efficiency

## Materials and Methods

### Strains and media

The fission yeast strain JB355 (h–, *fex1Δ fex2Δ his3-D1 leu1-32 ura4-D18 ade6-M216*) was used as the master strain for most transformations (available upon request), and replaces the strain JB224 we previously published [18] but which has mating defects. JB355 was created by transformation of FY527 using the standard lithium acetate procedure [21] and CRISPR-Cas9 method with the vector pJB174 targeting *fex1* and *fex2* (Cas9 cut site used : GGATAGGCCAGGCAAATGCA). The knockouts of *fex1* and *fex2* were confirmed by sequencing. We confirmed by sequencing that a single base is deleted in the cutting site used for editing, which leads to a frameshift for both genes. We sequenced this strain to confirm that no other mutation was present in its genome. All the strains created and used in this study are listed in Supplemental Table 1.

Yeast cells were grown in rich medium (YE5S; 0.5% yeast extract and 3% dextrose) supplemented with adenine, leucine, lysine, histidine and uracil (0.225 g/L each). Media preparation and basic manipulation methods of *S. pombe* were carried out as described previously [21].

*Escherichia coli* DH5α cells were used for subcloning polymerase chain reaction (PCR) products and were grown in LB medium.

### Vector preparation, design and cloning

The vector pJB166 (Addgene #86998) expressing SpCas9 under the adh1 promoter, the Fex1p selection marker under the ura4 promoter, and a sgRNA preceded by a *rrk1* leader and followed by a Hammerhead Ribozyme as described in [17] was digested and prepared for gap repair as follows: the vector pJB166 was purified from DH5α bacterial cells using the Genopure plasmid midi kit (Roche, 03143414001) and diluted to 1 μg/μL. 10 μg of plasmids were then digested overnight with the restriction enzyme CspCI (NEB, #R0645S) in a total volume of 50 μL. At this step, the complete digestion of the plasmid can be confirmed by running 1 μL of the digestion product on a 0.6 % agarose gel and observing a single band at about 11 kb. The digested plasmid was then purified using the QIAEXII Gel Extraction Kit (Qiagen, Q20021) and diluted to a final concentration of 100 ng/μL and ready for transformation.

The vector pJB174, targeting the genes *fex1* and *fex2* was amplified from pJB166 with the pairs of primers JB′330-JB’742 and JB′331-JB’743 and the 2 PCR products were annealed by Gibson assembly [22].

### Genome editing using the CRISPR/Cas9 and gap repair protocol

For genome editing with CRISPR/Cas9, we used the protocol described in [18] with the following modifications.. The strain JB355 (in which *fex1* and *fex2* are knocked out) was transformed using the lithium acetate method with 100 ng of pJB166 plasmid CspCI digested and purified, expressing Fex1p and Cas9; 1 μg of a PCR product containing annealing ends for pJB166 and a 20 bp Cas9 target specific sequence (see Figure 1B), and 1 μg of donor DNA, which contains a sequence with the intended edits flanked with sequences homologous to the genomic region around the Cas9 cut. The cells were resuspended in 1000 μL of water and we plated 150 μL on YE5S + 1 mM NaF and incubated at 32°C for 2–3 days. Twelve colonies were restreaked on YE5S + 1 mM NaF for a further 2–3 days. Colonies appearing after the restreak were directly screened using colony PCR.

**Figure 1:**
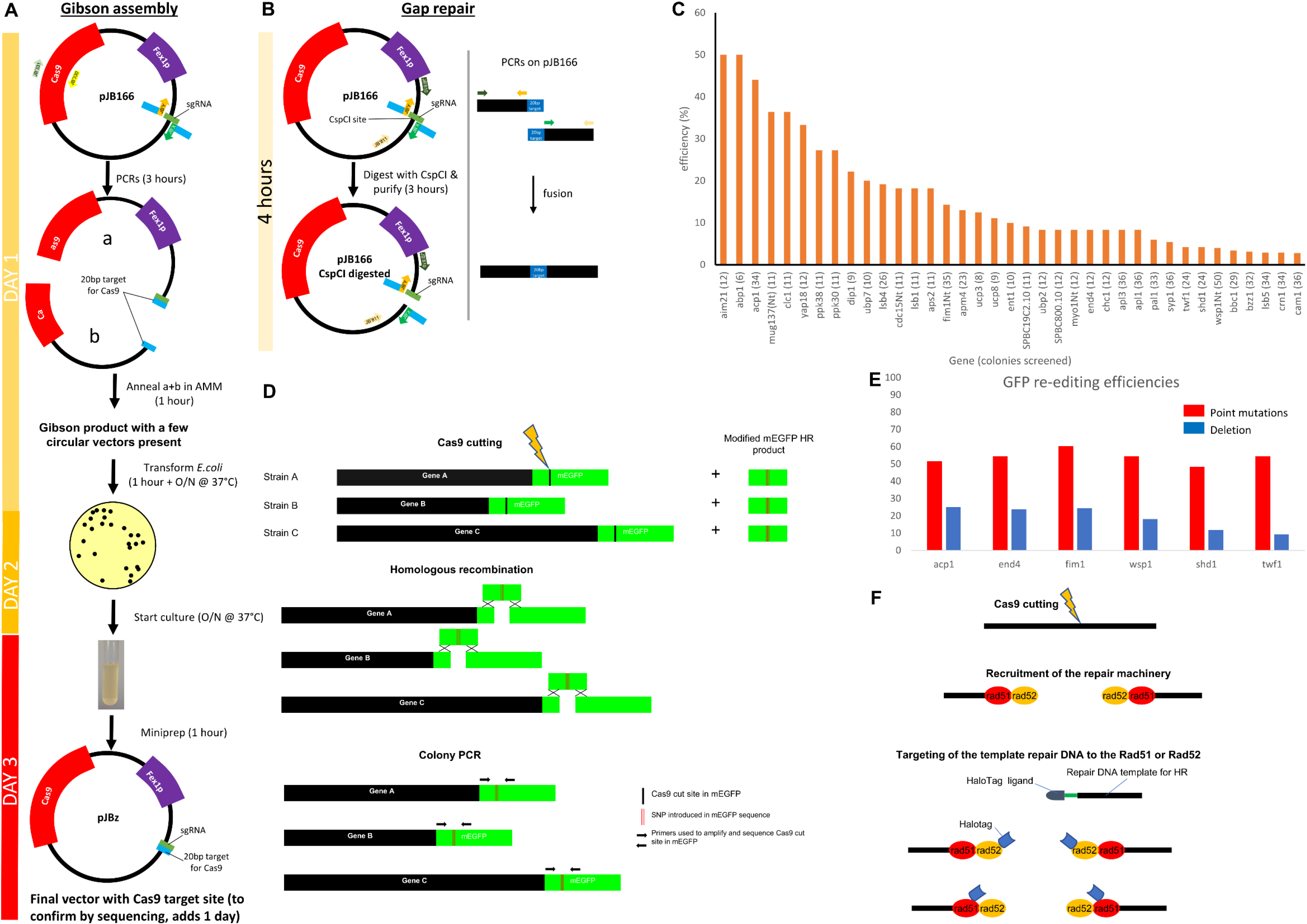
Fast CRISPR-Cas9 editing with gap repair demonstrates editing efficiency is not limited by homology search. (A) Traditional methods require the construction using Gibson assembly of the plasmid containing Cas9 and the gRNA and it purification, which takes about 3 days. (B) The gap repair method involves the digestion of pJB166 by CspCI and the purification of the digested vector. In parallel, the primers JB’X and JB’Y are used with respectively JB’811 and JB’810 and the two PCR products are annealed with PCR. The resulting PCR product can be directly transformed in yeast with the digested pJB166 vector and the two pieces are annealed *in vivo* by gap repair. It takes less than 4 hours to obtain the two products needed for the gap repair method. (C) Efficiency of gene editing is highly variable across the genome. Our library of mEGFP-tagged genes involved in endocytosis in *S. pombe* was created using the master strain JB355 (*fex1Δfex2Δ*), in which we used CRISPR/Cas9 editing with the gap repair method we present in this paper. The editing efficiency for each gene is reported and the number of colonies screened is indicated next to each gene name. Most genes were tagged in C-terminal, and we added “Nt” at the end of gene names that were tagged in N-terminal. (D) Strategy and comparison of editing efficiencies of the same DNA sequence at different locations in the genome. All strains (A, B, C, etc) have a single gene tagged with mEGFP. We induce a cut by Cas9 on the mEGFP DNA sequence at the same location in mEGFP in all strains using the same gRNA and donor DNA for HR repair. After Cas9 cutting and HR recombination, the mEGFP edited region is amplified by colony PCR and the number of colonies with the original or the edited sequences (mEGFP*) are determined by sequencing. (E) The efficiency of silent point mutation editing in mEGFP (red) is similar for all 6 strains ((average of 54%, Chi-square p-value=0.7, 79 to 93 colonies screened, 2 biological repeats for each strains). The efficiency of mEGFP deletion (blue) is also similar in all 6 strains (average of 18%, Chi-square p-value=0.3, 24 to 44 colonies screened, 2 biological repeats for each strains). (F) Schematic of our strategy to target the donor DNA to the Cas9 cut site. After Cas9 induces a double strand break, the 2 DNA ends are rapidly coated with DNA HR repair proteins. Our strategy was to modify the donor DNA so it carries a bromohexyl group at its 5’ end and to label two of the HR repair machinery proteins, Rad52p or Rad51p with a HaloTag, such that the donor DNA and one copy of Rad51p Rad52p can be covalently linked together.

### Colony PCR

After transformation, plating and restreak on YE5S + 1 mM NaF, colonies were screened using Q5 High Fidelity DNA Polymerase (NEB, M0491L) and at least one primer upstream or downstream of the DNA region used for recombination. PCR products were directly sent for sequencing (Keck DNA sequencing lab, New Haven, CT, USA).

### DNA Primers and Plasmids

All the primers used in this study are listed in Supplemental Table 2.

For the experiments where the donor DNA was targeted to the cut site, DNA primers were ordered from the Yale Keck Oligos Synthesis Resource and contained a 5’ bromohexyl group and Spacer 18 modifications.

All the plasmids used in this study are listed in Supplemental Table 3.

## Results

### CRISPR-Cas9 editing made faster and more efficient in fission yeast using gap repair to assemble the vector containing Cas9 and the gRNA

We have previously reported an improved protocol to edit the fission yeast genome using CRISPR-Cas9 [18]. The use of a fluoride channel as selection marker as opposed to auxotrophic markers usually used in the field significantly reduced the time from transformation to identification of mutants. At that point, one of the most time-consuming part of the protocol was the creation of the vector containing Cas9 and the gRNA. In our lab, the Gibson assembly method [22] was used to create these vectors, which contain a 20 bp sequence defining the Cas9 cutting site specific to each editing (Figure 1A). The cloning step was clearly the most time-consuming step, so to make our protocol as short and efficient as possible, we adapted this gap-repair strategy [20] to assemble the vector containing Cas9 and the Fex1p selection marker we developed [18], with the gRNA, as an alternative to cloning.

Our master plasmid pJB166, which contains everything but a specific sequence defining the gRNA, is digested with the restriction enzyme CspCI (NEB, #R0645S). This open vector is purified using the QIAEXII Gel Extraction Kit (Qiagen, Q20021) and can be stored at -20°C for up to 4 months (Figure 1A). The DNA fragment containing the sequence for the gRNA is assembled in two steps. The first step consists of creating two fragments with homology on each side of the CspCI site that also contain the DNA sequence encoding the gRNA or its reverse complement, and the second step consists of ligating these fragments via PCR. For the first step, we use universal primers (respectively JB’810 and JB’811), which have homology to sequences ∼350 bp upstream of the CspCI site on our master plasmid, and two specific primers (respectively JB’X and JB’Y), made of a variable 20 bp tail corresponding to the gRNA sequence or its reverse-complement and ∼20 bp homology with the vector sequence around the CspCI site. After obtaining the PCR products built with the JB’810-JB’Y and JB’X-JB’811 primer pairs, the tails added to JB’X and JB’Y allow the PCR amplification of a chimeric product carrying the 20 bp sequence for the gRNA target site flanked on each side with ∼350 bp sequences homologous to sequences of the master plasmid (Figure 1B). The transformation of both this construct and the digested master plasmid in fission yeast allow *in vivo* gap repair recombination [23] and the creation of the final gRNA vector in a faster manner than previously published protocols.

Typically, the preparation of the CspCI digested and column purified vector takes a few hours and yields enough product for up to 500 transformations. The limiting step is now the production of the chimeric PCR containing the 20 bp target site for Cas9. We typically create the chimeric DNA product on transformation day, in less than four hours, which renders experimental planning easier. This is a considerable improvement since the cloning of the 20 bp Cas9 cut site sequence using the Gibson assembly method takes at least 3 days (Figure 1A&B).

### Editing efficiency varies greatly along the fission yeast genome

While creating a library of mEGFP-tagged proteins involved in clathrin-mediated endocytosis (CME), we realized that the rate of positive colonies identified (i.e. with mEGFP inserted) varied a lot from gene to gene (Figure 1C). For each transformation, we assessed efficiency by first determining mEGFP insertion at the correct locus by PCR for 12 restreaked colonies, and in the event where we could not identify at least one positive, we screened an additional 24 colonies.

The genes *acp1, abp1, mug137, clc1, yap18* and *aim21* present the highest rates of editing with more than a third of the colonies screened showing mEGFP insertion with *abp1* and *aim21* even showing 50% of positives. Most genes present an efficiency of tagging between 5% and 27%, and some genes were harder to edit, with an efficiency of tagging under 5% (*crn1, wsp1, shd1, twf1, lsb5, bbc1, bzz1* and *cam1*). In some instances, after transformation and screening, we couldn’t identify any positives among the 36 colonies analyzed and had to design a new Cas9 cut site and perform a new transformation to obtain positives (e.g. *bbc1, ubp2, ubp7*).

It was recently suggested that insertion of the carrier DNA used during the transformation may occur [24] and hence may decrease the rate of positives. However, we have never observed such insertion in the hundreds of negative colonies we checked by PCR or sequencing. It is important to point out that the protocol used in [24] skips the denaturation of the carrier DNA usually performed when using the standard lithium acetate protocol [21], which may explain the unexpected insertions.

The editing variability we observed while creating our library led us to wonder what governed editing efficiency.

### CRISPR-Cas9 editing efficiency does not directly correlate with the genomic location of the target sequence

The CRISPR-Cas9 genome editing process in the experiments above involves two steps: a) the binding of Cas9 to its target site and subsequent cleavage, and b) the repair by homologous recombination at the severed site. Several factors could influence the efficiency of these two steps, including the physical location of the DNA region to edit. Epigenetic modifications such as methylation, acetylation or histone binding are widely but unequally spread along the genome and may impede the binding of Cas9 or proteins of the homologous recombination machinery [25-27]. Hence, we wondered whether Cas9 binding to its target site could be modulated by accessibility of the DNA at specific genomic locations and lead to the variabilities in editing efficiency we observed.

To study solely the effect of the genomic location on editing efficiency, we needed to be able to perform the exact same editing, i.e. using the same sequence for the Cas9 cut site and the same donor DNA used for the repair, but located at different positions on the genome (Figure 1D). For that, we took advantage of our library of strains, where different endocytic gene are tagged with mEGFP, which allowed us to target the same cut site within the mEGFP sequence and use the same DNA sequence for homology repair, but at different locations in the genome. We selected six strains (*acp1-mEGFP, end4-mEGFP, mEGFP-fim1, mEGFP-wsp1, shd1-mEGFP* and *twf1-mEGFP*) spanning a range of efficiencies from 4% to 44%. First, we edited mEGFP by introducing seven silent point mutations in the coding sequence using the same donor DNA for all strains. After transformation, colonies were screened by PCR with the same primers in each case, the PCR products were sequenced and the number of effectively edited colonies (i.e. carrying the induced point mutations) was determined (Figure 1E). Three independent transformations were performed for each strain and 48 colonies per transformation and per strain edited were sequenced. All six strains were edited with efficiencies of 54% on average, and the difference between strains was not statistically significant (Chi-square p-value=0.7) (Figure 3E, red bars).

Next, instead of introducing silent point mutations in mEGFP, we deleted the entire mEGFP sequence in each strain. Two independent transformations were performed for each strain and 24 colonies per transformation and per strain were screened for deletion. In this case, it is important to point out that another variable was introduced since the donor DNAs used by the HR machinery to delete mEGFP are different for each strain studied. In that case again, each strain presented a similar editing efficiency, but efficiencies were lower, 18% on average, and the difference between strains was not statistically significant (Chi-square p-value=0.3) (Figure 3E, blue bars).

Altogether, these results strongly suggest that the location of the chosen editing site on the genome doesn’t have a direct effect on the editing efficiency.

### Addressing the HR product to the Cas9 cut site fails to improve *lsb1* and *pil1* editing efficiencies

A key yet poorly understood step of HR is the homology search, where a nucleoprotein filament at the cut site searches for a corresponding homologous sequence to use as a template for repair [28-31]. We hypothesized that Cas9 genome editing could be limited by the efficiency of the homology search.

In an attempt to improve editing efficiency, we decided to create bioengineered cells to facilitate the homology search and the localization of the donor DNA to the cut site. To do so, we created an artificial bond between a protein that localizes to DNA double-stranded breaks and is involved in repair, and the donor DNA to be used as a matrix for cut repair by HR. We engineered two strains where Rad51p and Rad52p, two proteins involved in recombination and repair of damaged DNA by HR [32-35], have been fused to a HaloTag. In addition, we added to the donor DNA a 5’-bromohexyl group, which can covalently react with a HaloTag (Figure 1F). Using this strategy, we expected the homology search, and hence genome editing, to be more efficient. We first verified that the addition of a HaloTag to either Rad51p or Rad52p didn’t have a deleterious effect on the strain’s ability to perform HR. As a test editing, we tried to delete the *pil1* gene, since our lab has shown this editing is very efficient (around 50% in most conditions tested and up to 75%). The *HaloTag-rad51* (JB347) and *rad52-HaloTag* (JB364) strains did not show a significant decrease in *pil1* deletion efficiency compared to a wild type strain (45% and 75% vs 45% respectively, Table 1), so we felt confident both strains could be used for subsequent experiments. The addition of the 5’-bromohexyl group to the donor DNA used for *pil1* deletion didn’t significantly improve the efficiency of editing (55% vs 75% in JB347 and 67% vs 45% in JB364), compared to the same transformation in a wild-type strain where a HaloTag was not fused to Rad51p or Rad52p (64% vs 45%).

**Table 1:** Editing efficiencies when targeting the donor DNA to the cut site.

|  |  | Strain edited |  |  |
| --- | --- | --- | --- | --- |
| Editing | Ligand type on donor DNA | wt | HaloTag-rad51 | rad52-HaloTag |
| pil1Δ | No ligand | 45% | 75% | 45% |
|  | HaloTag | 64% | 55% | 67% |
| lsb1-mEGFP | No ligand | 8% | 4% | 13% |
|  | HaloTag | 19% | 4% | 8% |

Knowing that *pil1* deletion was already highly efficient, we didn’t expect our strategy to improve it much. The goal of this strategy was to improve the editing efficiencies of genes relatively hard to edit (e.g.: editing efficiency of 20% or less). We selected the *lsb1* gene as a good candidate (18% efficiency for mEGFP tagging, Figure 1C) and repeated that editing in a wild-type, HaloTag-rad51 and rad52-HaloTag strains, and with an HR product carrying or not carrying the 5’-bromohexylgroup. The transformations with HR products free of HaloTag ligand respectively yielded 8%, 4% and 13% of positive (24 colonies screened). The same transformations but with the 5’-bromohexyl modification on the donor DNA, making it able to conjugate with HaloTag on Rad51p or Rad52p produced respectively 19%, 4% and 8% of positives (24 colonies screened). Altogether, our data suggest that localizing the donor DNA to the HR repair machinery may not be a strategy to significantly improve CRISPR-Cas9 editing efficiency systematically in *S. pombe*.

## Discussion

Here we have demonstrated that the gap-repair method in combination with fluoride selection is an efficient strategy to perform fast genomic editing in fission yeast. According to our protocol, in which we use longer homology sequences between the two gapped products than previously reported, selection of correctly gap-repaired plasmids is not necessary to obtain high editing efficiency.

We also observed a wide variability in editing efficiency across the genome which appears not to be caused by the location of the editing site. Indeed, our results show that if the same specific sequence is targeted for Cas9 cutting and HR repair at different locations in the genome, the efficiencies are roughly the same. It is important to mention that we edited strains where the sequence for a mEGFP had already been inserted. Hence, it is possible that the introduction of this foreign DNA sequence triggered conserved similar epigenetic modification at all the sites tested, leading to the same editing efficiencies observed in Figure 1E. More factors regulating DNA accessibility, cutting and repair will need to be tested to identify what really defines and limits genome editing efficiency. We also noticed that introduction of point mutations is consistently highly efficient, around 50% (Figure 1E), which could be explained by the high homology between the donor DNA and the severed genomic sequence.

## Supporting information

Supplemental Table 1: List of S. pombe strains used in this study

Supplemental Table 2: List of primers used in this study

Supplemental Table 3: List of plasmids used in this study

## Supplemental Tables

**Supplemental Table 1: List of *S. pombe* strains used in this study**

**Supplemental Table 2: List of primers used in this study**

**Supplemental Table 3: List of plasmids used in this study**

